# AUTS2 expression within mammalian lineage: a predictor of neural networks involved in Autism Spectrum Disorders

**DOI:** 10.1101/2022.12.12.520025

**Authors:** Aude-Marie Lepagnol-Bestel, Yann Loe-Mie, Mounia Bensaid, Michel Simonneau

## Abstract

Autism susceptibility gene 2 (AUTS2) is a neurodevelopmental regulator associated with an autosomal dominant neurological syndrome with ASD-like (Autism Spectrum Disorder-like) features. The *AUTS2* syndrome phenotype includes borderline to severe intellectual disability (ID), microcephaly and mild dysmorphic traits. Specific ASD-like features, including obsessive or ritualistic behaviours, are frequently displayed, although sociability is largely unaffected. Syndrome severity is worse when mutations are present in the 3’ region (exons 9-19) of the *AUTS2* gene. *AUTS2* is also associated with alcohol consumption, heroin dependence, schizophrenia and dyslexia, using GWAS studies. Our working hypothesis is that *AUTS2* expression during mammalian evolution can be instrumental to predict phenotypes linked to the disease. For that, we studied brain In Situ Hybridization (ISH) data in mouse, marmoset, human embryo and fetuses and sequences data of Neanderthal, Denisovan and modern humans. Developmental mouse brain expression is found in neocortex, hippocampus, thalamus, tegmentum and cerebellum suggesting a link linked with cognition, stereotypies and perseverative behaviors. Expression in amygdala and claustrum found in marmoset extend these phenotypes to anxiety and higher-order cognitive processes. In human embryos (8 weeks) and fetuses (15- and 19-weeks), we found *AUTS2* expression in similar regions but also in medial, lateral and caudal ganglionic eminence, that are involved in the diversity of interneurons found in neocortex, hippocampus and striatum. Sequence comparison of *AUTS2* locus in Neanderthal, Denisova and modern human reveals novel sites in regulatory regions of *AUTS2*. Altogether, these results suggest that ISH distribution along mammals is a novel phenotype-related biomarker useful for translational research.

## INTRODUCTION

The autism susceptibility candidate 2 (AUTS2) gene on chromosome 7q11.22 at 7q11.2 was first identified and found disrupted as a result of a balanced translocation event (t7;20) ^1^ in a pair of monozygotic twins with autism spectrum disorder (ASD). Identification of more than 60 novel cases suggest that clinical phenotypes of AUTS2 related patients are more closely associated with intellectual disability (ID) rather than directly linked to classical features associated with ASD. Human *AUTS2* is a highly conserved gene that spans 1.2Mb. Human AUTS2 protein has two major isoforms, full-length (1259 aa) and C-terminal (711 aa). This short C-terminal protein is produced from an alternative transcription start site in exon. An AUTS2-Polycomb complex was shown to activate gene expression in the CNS ^2^.

Human *AUTS2* display nucleotide variants that define the human-Neanderthal sweep ^3^ and three human accelerated regions (HARs) ^4,5^.

Phenotypic analysis of AUTS2 syndrome patients was recently performed ^6^. All patients have borderline to severe ID/developmental delay, 83–100% have microcephaly. Mild dysmorphology are present. Behaviour is marked by it is a friendly outgoing social interaction. Specific traits of autism (like obsessive behaviour) are seen frequently (83%), but classical autism was not diagnosed in any.

In most of the cases, one can find a small in-frame deletions which are often inherited and give a mild clinical phenotype is associated with, Deletions and other mutations causing haploinsufficiency of the full length *AUTS2* transcript, that occur de novo, give a more severe phenotype and occur de novo. Previous studies have shown deletions within the C-terminus isoform spanning exons 9-19 are associated with a severe neurocognitive phenotype ^7^. AHDH was found in a patient was a partial duplication of 3’ part of *AUTS2* locus ^8^. ASDs have been described for full deletions and duplications of the *AUTS2* locus ^9^. Furthermore, diverse GWAS studies found that *AUTS2* locus is associated to a variety of neurological conditions such as addiction disorders ^10,11^, epilepsy ^12^, schizophrenia ^13,14^ and dyslexia ^15,16^.

The diversity of the disease manifestations of *AUTS2* variants within the brain underscore the importance of elucidating its role in neurodevelopment.

Our working hypothesis is that the analysis of the distribution of *AUTS2* transcripts during mammalian evolution can be instrumental to predict behavioral phenotypes in the different animal models used in neurobiology, in particular mouse and marmoset.

Here, we analyzed the distribution of AUTS2 transcripts in mouse, marmoset and human, during brain development and the evolution of AUTS2 locus by comparing sequences from Neanderthal, Denisovan and modern human.

## RESULTS

### Phylogenetic data from mouse to modern humans

To understand what phenotypes in animal models can be related to human AUTS2 syndrome ^6,7,17^, we took advantages of recent public databases of large-scale In Situ Hybridation (ISH) performed in two mammalian neurobiology models, mouse and marmoset. GenePaint is a digital atlas of gene expression patterns in various tissues and species with strong focus on mouse embryos (https://gp3.mpg.de/). Expression patterns are determined by non-radioactive *in situ* hybridization on serial tissue sections. Allen Brain Atlas is an anatomically comprehensive digital atlas containing the expression patterns of ∼20,000 genes in the adult mouse brain (www.mouse.brain-map.org). Recently, the open Marmoset Gene Atlas (https://gene-atlas.brainminds.jp/) established a genome-wide, high-resolution atlas of the gene expression throughout the common marmoset (Callithrix jacchus) brain. It uses in situ hybridization (ISH) analysis to systematically analyze changes in gene expression over the course of postnatal brain development to adult stage ^18,19^.

We also generated quantitative radioactive ISH data from human embryos as in ^20^. *AUTS2* was also shown to be implicated in human evolution, having several regions where its human sequence significantly changed when compared to Neanderthals and non-human primates. We used sequences from Neanderthal, Denisovan and modern human to analyze evolution of transcription factor binding sites in these regions.

Altogether, we cover the different branches, including mouse, marmoset, Neanderthal, Denisovan and modern human, that appeared from their ∼ 90 MY common ancestor (Fig.1).

**Figure 1.**
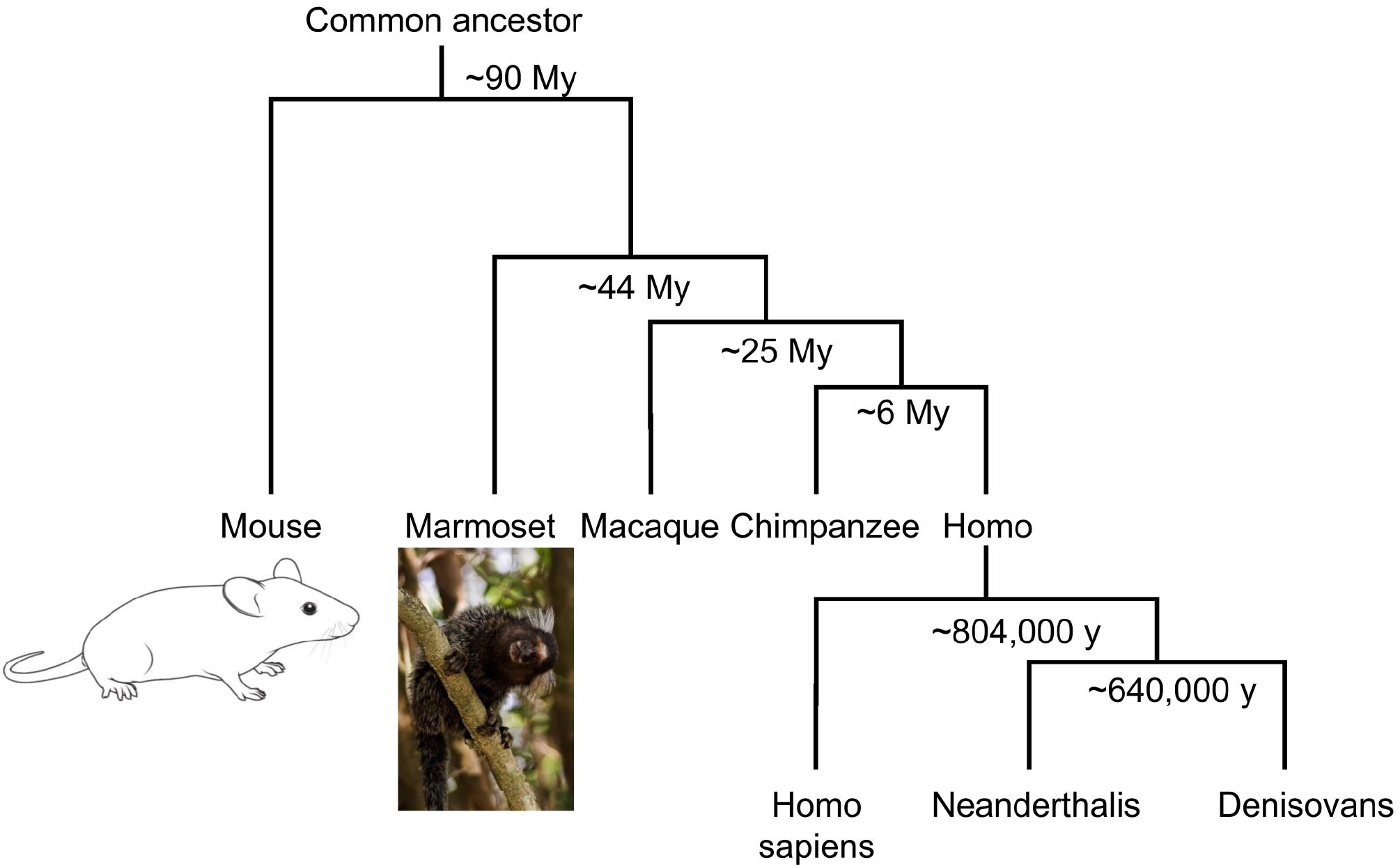
Phylogenetic tree relating the mouse, Marmoset *Denisova, Neanderthal* and Homo sapiens. DNA sequences of the Neanderthals and the Denisova were estimated to have diverged on average 640,000 years ago, and from present-day Africans around 804,000 years ago (Reich et al., 2010). Specimens are not drawn to scale.

### Auts2 expression in developing and adult mouse brain

We took advantage of public databases to analyze developing mouse brain (Genepaint (https://gp3.mpg.de/)) and adult mouse brain (Allen Brain Atlas at www.mouse.brain-map.org).). *Auts2* expression is observed in multiple areas of E14.5 mouse brain, including neocortex, hippocampus, dorsal thalamus, Septum, Striatum, Olfactory epithelium, Hypothalamus, Tegmentum, cerebellum and medulla oblongata (Fig. 2).

**Figure 2.**
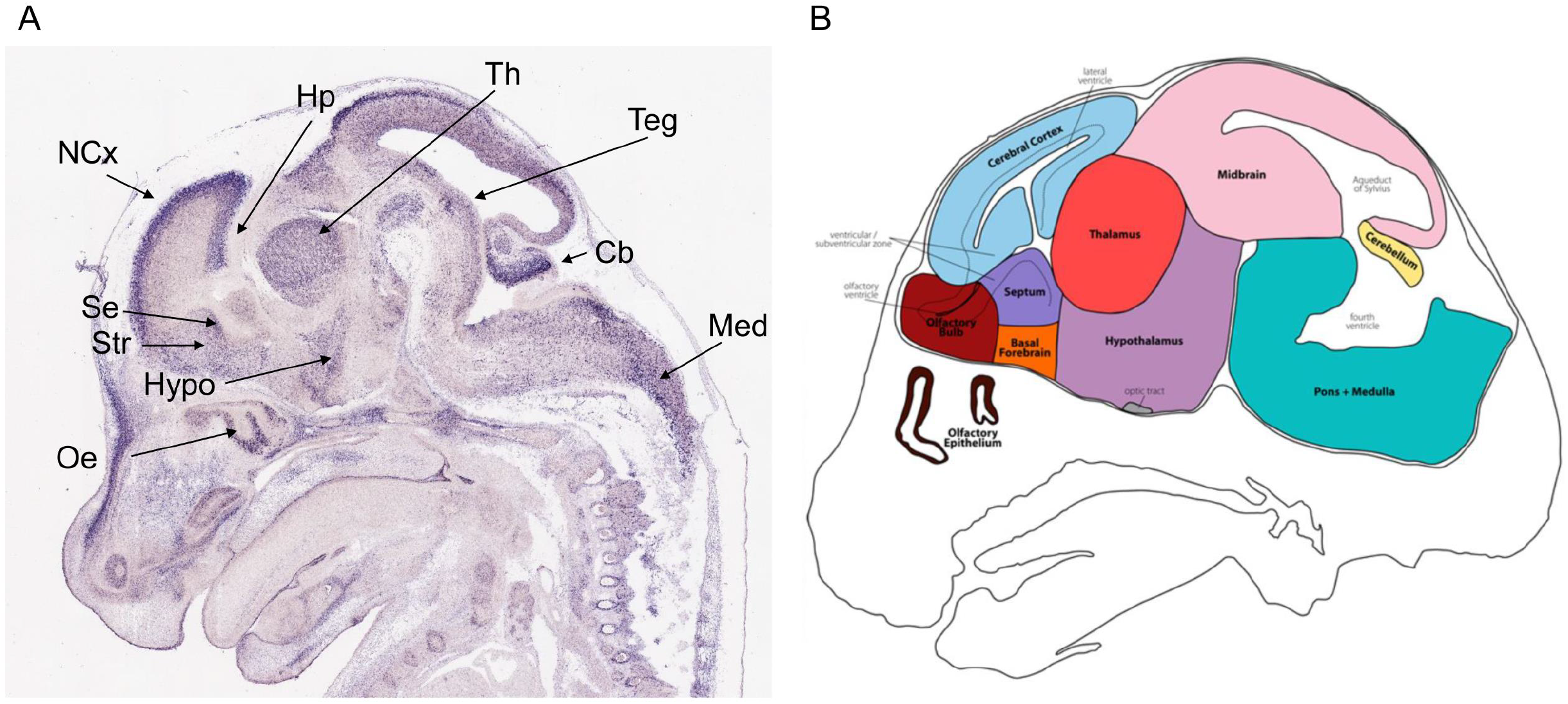
*AUTS2* expression in developing mouse brain. **A**. By *in situ* hybridization, *Auts2* expression is observed in multiple areas of E14.5 mouse brain, including Neocortex (Ncx), hippocampus (Hp), dorsal thalamus (Th), Septum (Se), Striatum (St), Olfactory epithelium (Oe), Hypothalamus (Hypo), Tegmentum (Teg), cerebellum (Cb), and medulla (Med). **B**. Schematic of main brain regions for E14.5 mouse (from GeneSat). Sagittal section from Genepaint (https://gp3.mpg.de/).

Interestingly, concerning the expression of *Auts2* in the tegmentum, it was demonstrated that virtually all of the dopaminergic TH positive neurons, that are located in tegmentum, in particular in Subtantia Nigra (SN) and Ventral Tegmental area (VTA) express *Auts2* ^21^. Interestingly, SN and VTA are located at the posterior region of the tegmentum, at the junction between midbrain and pons (Fig. 2B).

By *in situ* hybridization, *Auts2* expression is observed in multiple areas of the adult brain (C57BL/6J strain; 56 days of age; male) on a coronal section (Fig.3A). Higher expression is found in Dentate Gyrus (DG), Cornu Ammonis 3 (CA3), subiculum, lateral entorhinal cortex and temporal association areas. Visual area, Auditory Cortex and Cortical amygdalar area, posterior part medial zone express *Auts2* (Fig. 3A). Furthermore, in adult mouse cerebellum (sagittal section), *Auts2* transcripts localized in the layer of Purkinje cells (Fig. 3B). In the E14.5 mouse brain, *Auts2* expression was homogeneous along the rostro-caudal axis. However, by E16, the Auts2 expression pattern was reported to change with changed with an expression limited to superficial layers ^21^.

**Figure 3.**
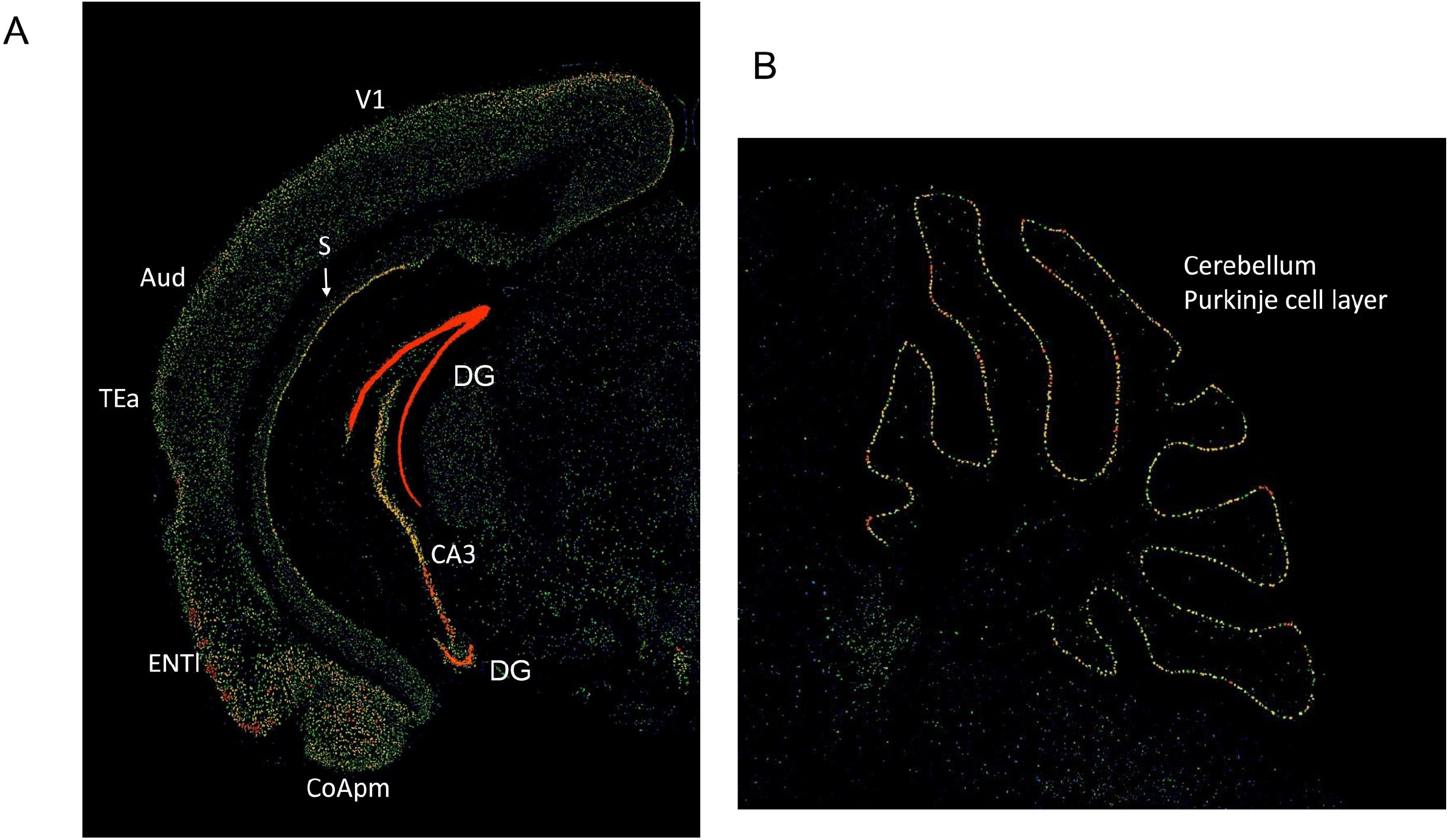
*AUTS2* expression in adult mouse brain. **A**. By *in situ* hybridization, *Auts2* expression is observed in multiple areas of the adult brain; on a coronal section. Higher expression is found in Dentate Gyrus (DG), Cornu Ammonis 3 (CA3), Subiculum (Sub), Visual area (VI), Auditory Cortex (Aud), Temporal association areas (TEa), Entorhinal area, lateral part (ENTl) and Cortical amygdalar area, posterior part medial zone (COApm). **B**. In adult mouse cerebellum (sagittal section), Auts2 transcripts localized in the layer of Purkinje cells. Data from Allen Brain Atlas.

These patterns of expression are in full agreement on previous report of *Auts2* expression ^21,22^. From these patterns of expression, it is possible to correlate Auts2-expressing neuronal networks with phenotypes related to ASDs, IDs, alcohol consumption or dyslexia. Hippocampus, septum, hypothalamus and cerebellum are known to be involved on social communication ^23^.Prefrontal cortex and striatum are implicated in stereotypies and perseverative behaviors ^24^. Prefrontal cortex, DG, CA3, subiculum and lateral entorhinal cortex participate to memory and cognition ^25^. Prefrontal cortex, striatum, SN and VTA are part of the reward circuit linked to alcohol consumption ^26^. DG, CA3, subiculum and lateral entorhinal cortex are involved in recognition memory that is linked to dyslexia ^27,28^.

### *AUTS2* expression results in the developing and adult non-human primate brain: the marmoset

Recently, the open Marmoset Gene Atlas (https://gene-atlas.brainminds.jp/) has published AUTS2 expression results in the developing and adult non-human primate brain. In the marmoset neonatal brain, *AUTS2* expression can be visualized in the Prefrontal cortex, in particular in Brodmann area 24 (A24; Anterior Cingulate Cortex), A6 and A8 (Fig. 4A-B) and in movement-control related areas (Caudate, Putamen,Thalamus) (Fig. 4A-B; Fig. 4C). Other regions with high *AUTS2* expression are cortical regions receiving sensory afferences: Piriform cortex, Orbital Proisocortex (Opro) and Gustatory cortex (Fig. 4A). Interestingly, *AUTS2* is also express in the claustrum (Fig. 4A).

**Figure 4:**
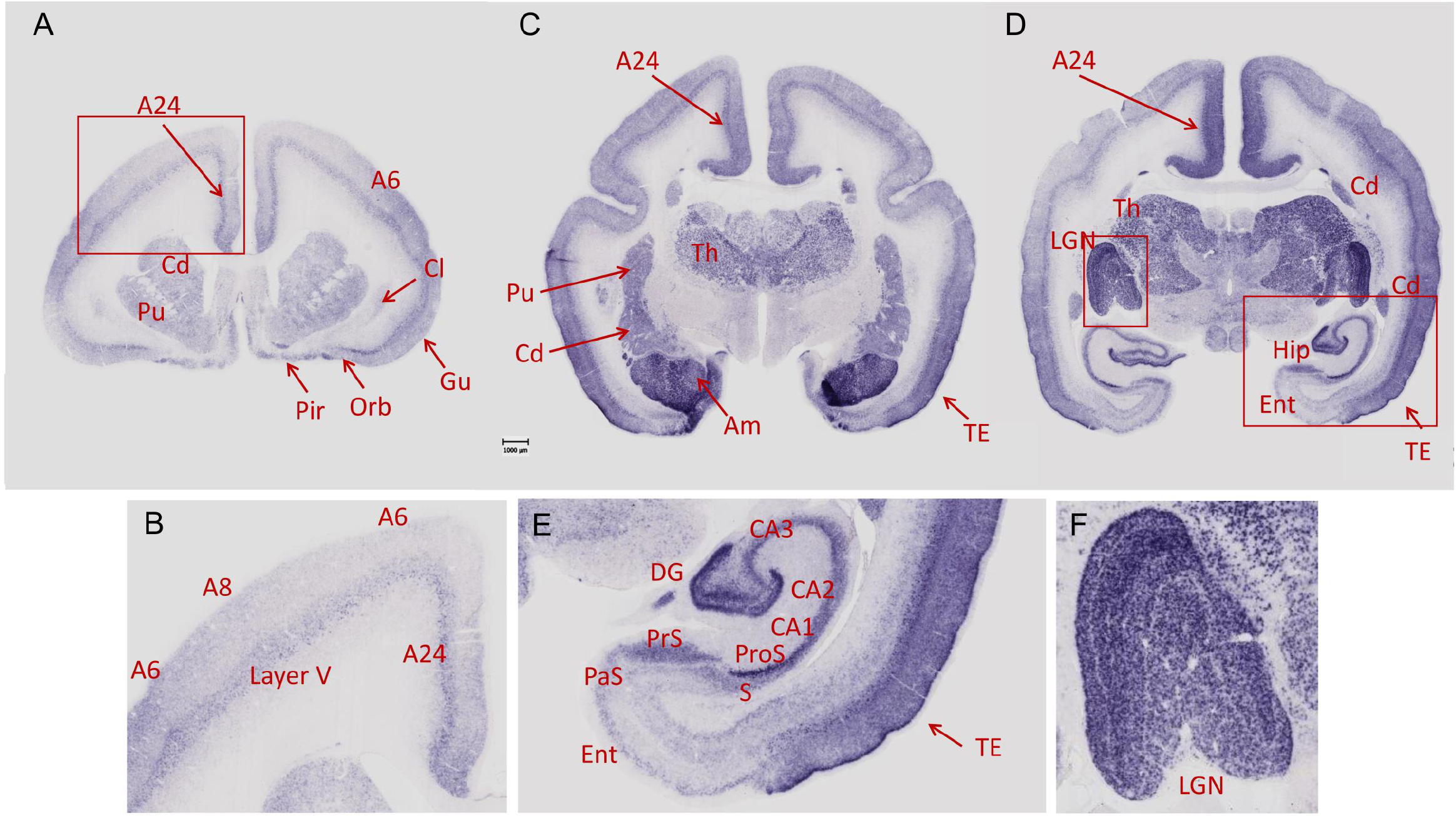
*AUTS2* expression in neonate marmoset brain. **A, C** and **D** sections are coronal sections from anterior to posterior axis. **A**. The highest levels of AUST2 are detected in Brodmann A24 (anterior cingulate cortex), caudate (Cd), putamen (Pu), claustrum (Cl), Brodmann A24 (anterior cingulate cortex), A6 and A8 that are all part of the frontal cortex, Piriform cortex (Pir), Orbital Proisocortex (Opro) and Gustatory cortex (Gu). **B**. Enlargement of the Brodmann A24 (anterior cingulate cortex) (red frame in **A**). Expression of AUTS2 is found in Brodmann A24 (anterior cingulate cortex), Brodmann A6 and A8, that all are part of the frontal cortex. Note that expression is higher in layers V for the Brodmann A24 (anterior cingulate cortex) and Brodmann A6, A8 areas. **C**. The highest levels of AUST2 are detected in Brodmann A24 (anterior cingulate cortex), Thalamus (Th), caudate (Cd), putamen (Pu), Amygdala (Am) and Temporal cortex area (TE). **D**. The highest levels of AUST2 are detected in Brodmann A24 (anterior cingulate cortex), Temporal cortex (TE), caudate (Ca), Thalamus, Hippocampus and related structures and Lateral Geniculate Nucleus (LGN). **E**. Enlargement of the Hippocampus and related structures (red frame in **D**). Hippocampus: Dentate Gyrus (DG), Cornu Ammonis 1 (CA1), Cornu Ammonis 2 (CA2), Cornu Ammonis 3 (CA3), Subiculum (S) Presubiculum (PrS), Prosubiculum (ProS) and Parasubiculum (PaS). Entorhinal cortex (Ent) and Temporal cortex area (TE). Note that expression is higher in layer V of the TE. **F**. Enlargement of the LGN region (red frame in **D**). Note that expression of AUTS2 is found in all layers of LGN (magnocellular, parvocellular and koniocellular layers). Data from the Marmoset Gene Atlas (https://gene-atlas.brainminds.riken.jp/).

Hippocampus and related regions including lateral entorhinal cortex and Temporal cortex express *AUTS2* (Fig. 4C; Fig 4E). High levels of AUTS2 are also found in amygdala (Fig. 4C-E) and in all layers of Lateral Geniculate Nucleus (LGN) (Fig. 4D; Fig. 4F).

Hippocampus and related regions can be involved in memory defects. Interestingly, CA2 that is known as a critical hub of sociocognitive memory processing ^29,30^, highly expressed *AUTS2*. Fronto-striatal pathways are involved in stereotypies and perseverative behaviors ^24^. Expression in Amygdala. suggests possible implication in anxiety and fear associative memory ^26,31^. The claustrum is a brain region that has been investigated for over 200 years but its precise function remains unknown ^32^. Sir Francis Crick with Christof Koch suggested that the claustrum can be critically linked to consciousness ^33^. Widespread extensive connectivity of single claustrum neurons with the entire cerebral cortex suggests a prominent role in higher order processes ^34^.

Expression in LGN can be related to the involvement of the visual magnocellular pathway in ASDs ^35,36^.

In adult marmosets, *AUTS2* mRNA levels remain high only in the amygdala, and in hippocampal dentate gyrus.

### AUTS2 expression analyzed in the developing human brain

We analyzed expression of *AUTS2* at three stages of human brain development: 8, 15, 18 and 22 weeks respectively (Fig. 5; Fig.6). We used radioactive antisense riboprobes. *AUTS2* expression was quantified by optical imaging of the spatial distribution of beta-particles emerging from brain sections ^20,37^.

**Figure 5:**
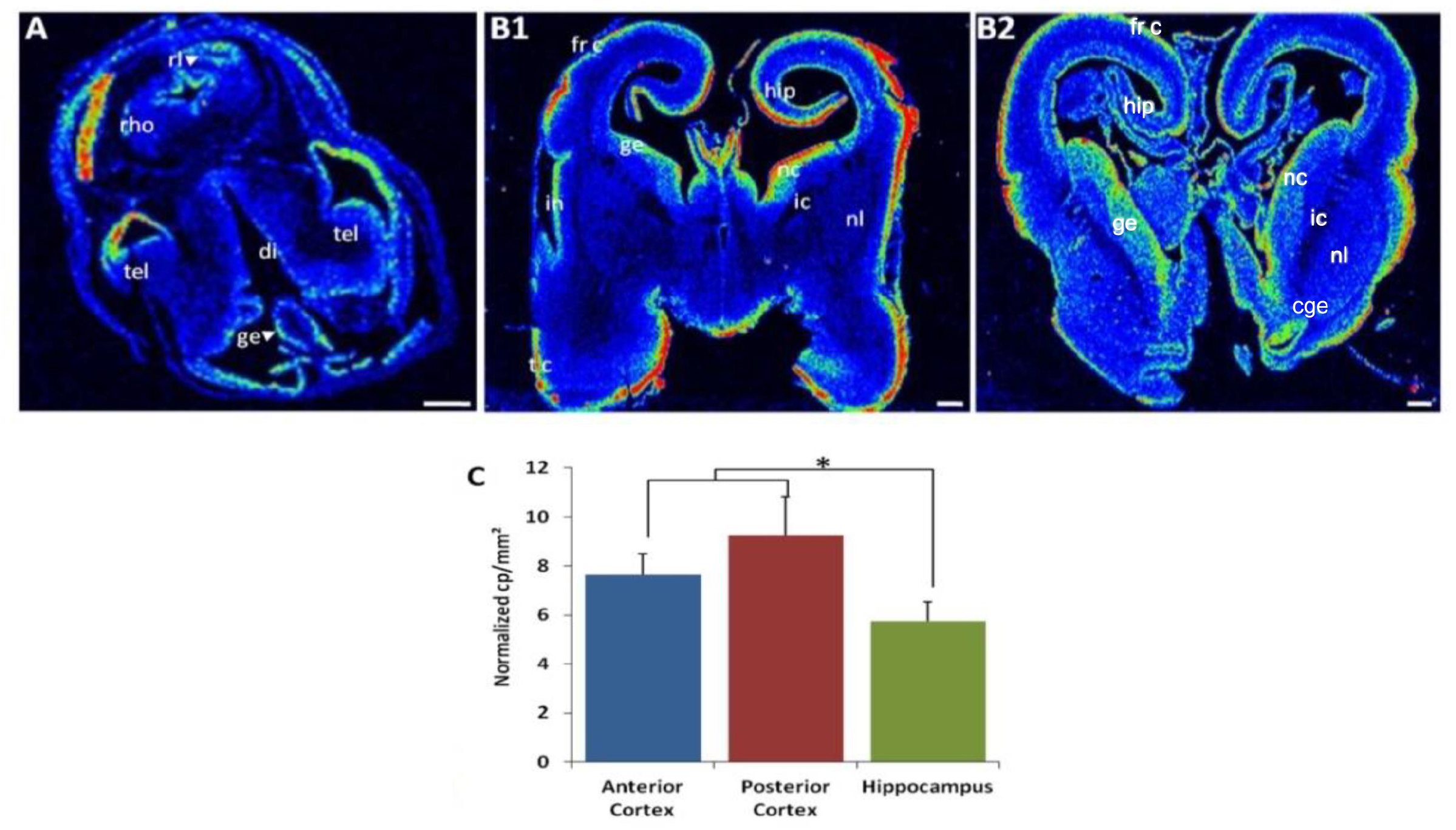
AUTS2 expression during human brain development. **A-B**. Coronal sections of 8-week (A) and 15-week-old (B) human embryos (B1 anterior section and B2 posterior section) hybridized with *AUTS2* antisense radioactive riboprobe. di: diencephalon; ic: internal capsule; fr c: frontal cortex; ge: germinal zone; hip: hippocampus; in: insular cortex; mes: mesencephalon; nc: nucleus caudate; nl: nucleus lenticular; rho: rhombencephalon; rl: rhombic lip; tc: temporal cortex; tel: telencephalon. Scale bar=1mm. **C**. quantification indicating expression of *AUTS2* both in cortex and hippocampus at 15-week-old human embryonic development with a significantly higher expression level in cortical regions than in hippocampus. *

**Figure. 6:**
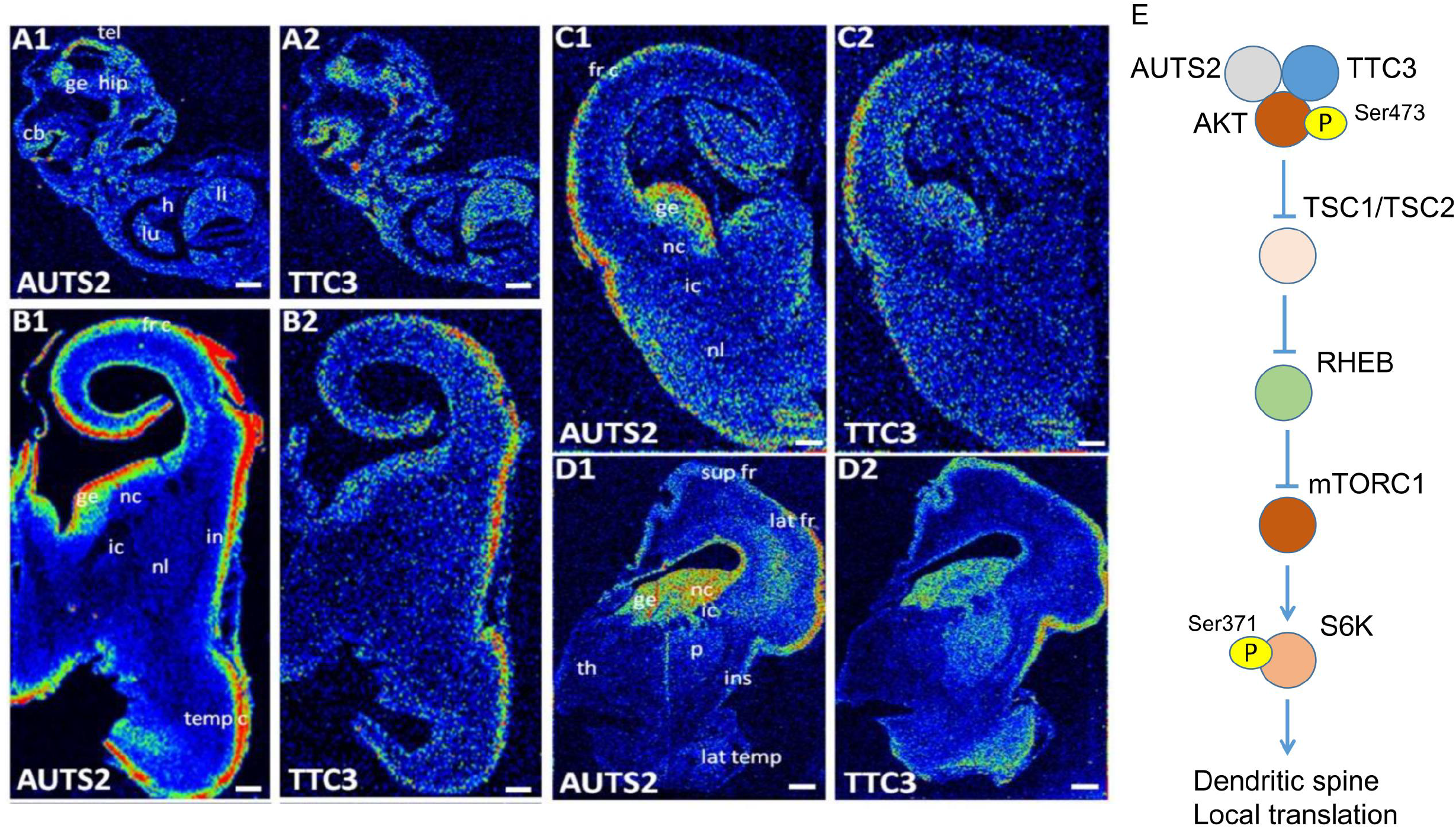
AUTS2 and TTC3 mRNA levels in the human central nervous system during early development and at mid-gestation. **A**. Sagittal sections of 8-week-old human embryos hybridized with AUTS2 (A1) and TTC3 (A2) antisense radioactive riboprobes. The two transcripts were detected in telencephalon (tel), ganglionic eminence (ge), hippocampus anlagen (hipp), cerebellum anlagen (Cb) and liver (li) but not in heart (h) and lung (lu). **B-C**. Coronal sections of 15-week-old human brains hybridized with AUTS2 (B1-C1) and TTC3 (B2-C2) antisense radioactive riboprobes. The two transcripts were detected in frontal cortex (fr c), temporal cortex (temp c), insular cortex (in) and germinal zone (ge) but nor in nucleus caudate (nc), nucleus lenticular (nl) and internal capsule (ic). **D**. Coronal sections of 19-week-old human brains hybridized with AUTS2 (D1) and TTC3 (D2) antisense radioactive riboprobes. The two transcripts were detected in frontal superior cortex (sup fr), lateral frontal cortex (lat fr), insular cortex (ins), ganglionic eminence (ge), internal capsule (ic) and nucleus caudate (nc) but not in thalamus (th), putamen (p) and lateral temporal cortex (lat temp). Scale bars=1mm. **E**. mTORC1 pathway that involves AUTS2-TTC3-AKT complex (modified from Lepagnol-Bestel et al., 2022).

In 8-week human embryo (Fig 5A), *AUTS2* is expressed in rhombencephalon; rhombic lip and germinal eminence. Ganglionic eminences (GE) are subcortical structures of gray matter which appear during the 5^th^ week post-fertilization on the floor of telencephalic vesicles ^38^.

In 15-week-old human embryos (Fig 5B; B1 anterior section and B2 posterior section), *AUTS2* is expressed in frontal cortex, hippocampus, temporal cortex, insula and ganglionic eminence (GE). At this stage, both MGE and LGE ganglionic eminence express *AUTS2* as visualized on an anterior coronal (frontal) section in Fig. 5B1. On a posterior coronal section (Fig. 5B2), AUST2 expression can be visualized in medial (MGE), lateral (LGE) and caudal (CGE)

The GE is anatomically subdivided into the MGE, the LGE and the CGE. These transient structures generate main neuronal networks. The MGE and LGE develop into the basal ganglia, striatum and pallidum respectively ^39,40^. The LGE generates projection neurons to the striatum, the medium spiny neurons that form 90% of the neuronal striatum population. The CGE gives rise to amygdala ^39,40^. Ganglionic eminences also generate a variety of interneurons. From the LGE, interneurons migrate to the olfactory bulb. The CGE produces interneurons migrating to the cerebral cortex ^41,42^. The MGE is a main source of interneurons throughout the cortex, hippocampus and striatum after tangential migration ^43,44^.

Quantification of *AUTS2* expression both in cortex and hippocampus at 22-week-old human embryonic development with a significantly higher expression level in cortical regions than in hippocampus (p<0.05) (Fig. 5C).

We recently demonstrated that AUTS2 directly interacts with TTC3 a E3 ligase of AKT that regulates dendritic spine function via mTORC1-dependent local translation ^45^ (Fig. 6E). Brain structures that co-express AUTS2 and TTC3 are expected to display modified organization of their neurons or of their neuronal networks.

In 8-week-old human embryos hybridized with AUTS2 and TTC3 antisense radioactive riboprobes, we found the two transcripts in telencephalon, ganglionic eminence, hippocampus anlagen and cerebellum anlagen (Fig. 6A1-A2). In 15-week-old human brains, AUTS2 and TT3 are co-expressed in frontal cortex, temporal cortex, insular cortex and germinal zone but nor in nucleus caudate (nc), nucleus lenticular (nl) and internal capsule (ic) (Fig. B1-B2, Fig. C1-C2). In 19-week-old human brains AUTS2 and TT3 are co-expressed in frontal superior cortex, lateral frontal cortex, insular cortex, ganglionic eminence, internal capsule and nucleus caudate (Fig. D1-D2).

From these patterns of co-expression, one expect to have phenotype changes in cognition (frontal cortex and cortical interneurons from MGE), in learning & memory (Hippocampus anlagen and hippocampal interneurons from GE), in social brain (Frontal cortex, insula and cerebellum) and stereotypies and perseverative behaviors (caudate, striatum interneurons from GE).

### A human-accelerated evolution of AUTS2 locus

Human AUTS2 and its mouse homolog encodes highly conserved protein with 85% of conservation in 1259 and 1261 amino acid sequences, respectively.

AUTS2 region 2 (377,373bp) was identified as displaying a signature of positive selection in the human compared to Neanderthal lineage 14 (Figure 7A-B). The number of mutations (9,207 in region 2) is statistically different (p=0.02) compared to region 1+3 (20,981 in 835,211bp). We analyzed the putative transcription factor sites that can be statistically different between the Human and Neanderthal AUTS2 locus. We found that 10 and 8 putative sites that are statistically significant (Zscores ≥3 or ≤3) for Human and Neanderthal AUTS2 loci. Human-associated TF sites are disproportionately associated with neuronal expression using Entrez Gene neuronal annotations (p=2.51x10 -4; Fisher’s exact test) (Figure 7C). These results suggest that Human AUTS2 gene displays a neuronal regulation of its expression contrasted in comparison to Neanderthal AUTS2 locus.

**Figure 7:**
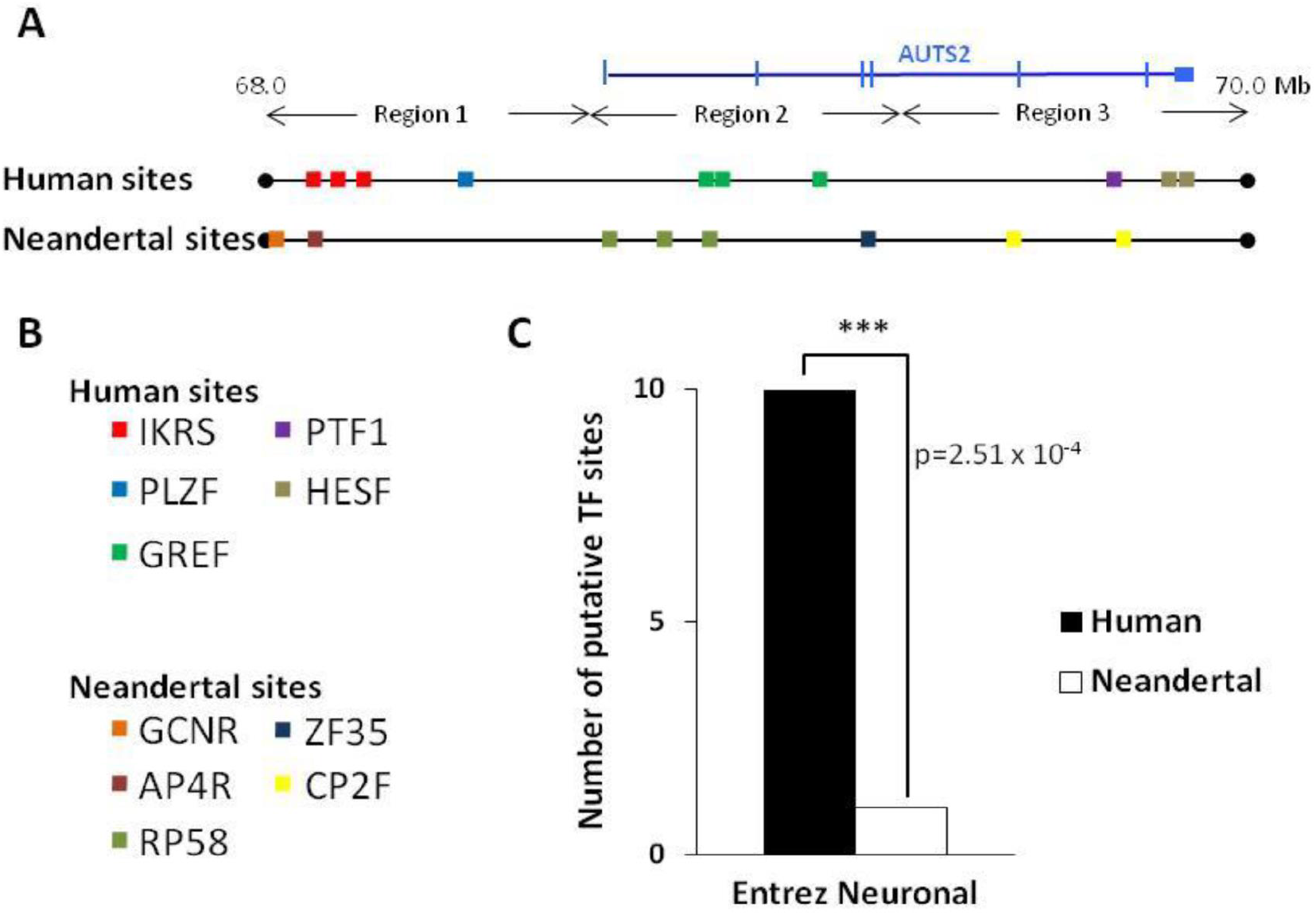
Human *AUTS2* gene displays a positive selection compared to Neanderthal lineage. **A**. Diagram of the chromosome 7 locus including the *AUTS2* gene. Region 2 (377,373bp) was identified as displaying a signature of positive selection in the human compared to Neandertal lineage (Green et al., 2010). Statistically significant (Zscores ≥3 or ≤3) putative transcription factor (TF) matrix sites using Genomatix suite are indicated for Human and Neanderthal AUTS2 loci. **B**. Indication of the TF matrix family name for the Human and Neanderthal sites. **C**. Human-associated TF sites are disproportionately associated with neuronal expression using Entrez Gene neuronal annotations (p=2.51x10^−4^; Fisher’s exact test).

These results suggest that Human AUTS2 gene may display a more complex spatial and temporal pattern neuronal regulation of its expression when compared to the Neanderthal AUTS2 locus. Moreover the observed differences in transcription factor sites at the AUTS2 locus may signify a novel expression pattern of the human as compared to Neanderthal AUTS2 locus, raising the possibility that these changes may have contributed to the brain evolution during the ∼400,000 years separating Neanderthal and Human lineages.

## DISCUSSION

Recent identification of risk genes for psychiatric disorders have set the stage for functional interrogation of disease related-circuits and underlying mechanisms of pathophysiology ^46^. Here, we propose that *AUTS2* expression within mammalian lineage can be a predictor of neural networks involved in Autism Spectrum Disorders. Our working hypothesis is that the distribution of *AUTS2* transcripts in different mammalian models can be instrumental to predict behavioral phenotypes.

Behavioral studies have been conducted using several different types of *Auts2* mutant mice. ^2,22,45,47–49^.

In the first study, across early development, KO mice were deficient in ultrasonic vocalizations emitted ^2^. In the study of Hori et al., 2015, *Auts2* heterozygotic mutant mice displayed behavioral abnormalities in anxiety-related emotions (elevated plus maze; cued fear associative memory but no changes in contextual fear associative memory) and recognition memory abnormalities.

Three studies were based on conditional knockout models leading to knockout of *Auts2* in excitatory neurons of the forebrain ^48^, in the cerebellum ^22^ or in developing forebrain ^49^. Selective deletion of *Auts2* in excitatory neurons in the adult forebrain ^48^ induces a phenotype different from the constitutive knockout ^47^. In this conditional knockout, KO mice display social deficits and altered vocal communication. Interestingly, the number of ultrasonic vocalizations (USVs) but also the complexity of USV syllables emitted from male mice during courtship behaviors are reduced ^48^. Selective deletion of *Auts2* in the cerebellum induces behavioral impairments in motor learning and vocal communications ^22^. Selective deletion of *Auts2* in developing forebrain led to hypoplasia of Dentate Gyrus with social deficits and stereotypies (excessive grooming and digging behaviors) ^49^.

The third type of *Auts2* mouse model developed by our group ^45^ is based on either deletion or duplication of the ∼1.2 Mb *Auts2* locus. We evidenced repetitive and restricted behaviors (rearing) in Del/+ mice.

Auts2 gene expression was found increased by repeated cocaine administration specifically in D_2_-type medium spiny neurons in the nucleus accumbens ^50^.

Altogether, these results indicate that Auts2 is involved in cognition and recognition memory, social memory, stereotypies and perseverative behaviors, anxiety and addiction.

Cognition is related with Auts2 expression in frontal cortex and hippocampus. Defects in Auts2 in these regions can be linked to intellectual disability (ID) found in patients.

Recognition memory involves hippocampus and related structures such as Lateral Entorhinal Cortex that express AUTS2 in developing marmoset brain. Recognition memory circuitry is linked to dyslexia ^27,28^. AUTS2 locus was recently found associated with dyslexia ^16^. Social brain involves frontal cortex, temporal cortex and amygdala ^51^ but also hippocampus and hypothalamus ^23^. Deregulation of Aust2 expression in these regions can induces abnormal social interactions and altered vocal communication as found in the *Auts2* mouse models. Stereotypes and perseverative behaviors (excessive grooming and digging behaviors; rearing) are linked to frontal cortex-striatum-thalamus loops that express *Auts2* in mouse and marmoset. Anxiety changes were evidenced in different models suggesting an expression of *Auts2* in amygdala. This expression is well documented in neonate marmoset.

AUTS2 expression is found in claustrum, in neonate marmoset. This region is linked to high-level cognition ^32^. Interestingly, each single claustrum neuron in mouse brain displays widespread extensive connectivity with the entire cerebral cortex ^34^.

Further work is needed to demonstrate if Aust2 is expressed in claustrum neurons in mouse. We also found AUTS2 expression in ganglionic eminence (GE) in human fetal brain. EMINENCE Subcortical structures such as striatum, pallidum and amygdala are derived from progenitors that originate in GE ^40,52^. GE generate interneurons whose number and diversity increase in primates compared to rodents ^53,54^. It would be important to identify if mouse GE and/or interneurons derived from GE express Auts2 and to manipulate their level of Auts2 expression.

Our data can permit to design future studies in selecting ether mouse or marmoset transgenic models to study *Auts2* expression impacts in subset of interneurons.

Altogether, these results suggest that ISH distribution along mammals is a novel phenotype-related biomarker useful for translational research.

## MATERIAL & METHODS

### Human sample preparation

Tissues were obtained from spontaneously or voluntarily terminated pregnancies following the informed consent of the parents and according to the French Ethical Committee recommendations. We studied embryos of 8 weeks and fetuses of 15 and 19 weeks. Tissues were fixed in 4% paraformaldehyde, embedded in paraffin and sectioned (5 μm).

### Probe synthesis and in situ hybridization

We used human cDNA clones from the RZPD Library (ID DKFZp547C245Q2; ID IRALp962D1923Q2). We synthesized ^35^S-labeled riboprobes for Nogo (700 bp riboprobe) and NgR (1200 bp), using the P1460 riboprobe in vitro transcription system (Promega). Hybridization was carried out as previously described (Lepagnol et al., 2008). Hybridizations were performed with both antisense and sense riboprobes. No signal was obtained with sense riboprobes. Expression was quantified with a Biospace Micro Imager and Betavision analysis software (Biospace Instruments) (<u>Charpak et al., 1989</u>). Additional adjacent sections were stained with hematoxylin/eosin/safranin (HES) for histological examination.

### Transcription factor sites analysis

Neanderthal sequences were downloaded from “UCSC Neanderthal portal”. We used Neanderthal contigs (mixed of 6 specimens sequenced). We selected fragments aligned on hg18 reference and analyzed only human sequences which had Neanderthal homologues. The analysis was conducted on sequences having exactly the same number of human and Neanderthal nucleotides. We obtained 12,437 pairs containing 1,212,584 nt for each species. We excluded from the analysis pairs which were identical between human and Neanderthal. This procedure reduced the dataset to 7,193 sequence pairs containing 797,226 nt (r1 254,125 nt, r2 241,463 nt, r3 301,638 nt) AUTS2 locus was analyzed with the genomatix matinspector program for transcription factor sites discovery with stringent conditions (core sim = 0.95 and opt +0.1) for both species. We counted only site predictions which were present in one lineage on sequence pair. For comparison, we randomly selected aligned regions (200,000 pair containing 18,890,929 nt, with elimination of identical pairs it was reduced to 115,069 pairs containing 12,312,338 nt) across whole genome and did the same analysis. We used boostrap approach to infer significance of matrix family hits on AUTS2 locus. We randomly chose fragment pairs to obtain, at least, the number of nucleotides of the region in the randomly selected regions, counted the number of matrix family hits and did this analysis 10,000 times. We could estimate the number of matrix family hits for region of approximately the same length of the region of interest and calculated z-score for each matrix family and by region.

## ACKNOWLEDGMENTS

This work was founded by an European Grant ERA-Net Cofund Action on Nanomedicine under Horizon 2020 Euronanomed 3 (project MoDiaNo) to M.S.

## References

1. Sultana, R., Yu, C.-E., Yu, J., Munson, J., Chen, D., Hua, W., Estes, A., Cortes, F., de la Barra, F., Yu, D., et al. (2002). Identification of a novel gene on chromosome 7q11.2 interrupted by a translocation breakpoint in a pair of autistic twins. Genomics 80, 129–134. 10.1006/geno.2002.6810.

2. Gao, Z., Lee, P., Stafford, J.M., von Schimmelmann, M., Schaefer, A., and Reinberg, D. (2014). An AUTS2-Polycomb complex activates gene expression in the CNS. Nature 516, 349–354. 10.1038/nature13921.

3. Green, R.E., Krause, J., Briggs, A.W., Maricic, T., Stenzel, U., Kircher, M., Patterson, N., Li, H., Zhai, W., Fritz, M.H.-Y., et al. (2010). A draft sequence of the Neandertal genome. Science 328, 710– 722. 10.1126/science.1188021.

4. Oksenberg, N., Stevison, L., Wall, J.D., and Ahituv, N. (2013). Function and regulation of AUTS2, a gene implicated in autism and human evolution. PLoS Genet 9, e1003221. 10.1371/journal.pgen.1003221.

5. Whalen, S., and Pollard, K.S. (2022). Enhancer Function and Evolutionary Roles of Human Accelerated Regions. Annu Rev Genet 56, 423–439. 10.1146/annurev-genet-071819-103933.

6. Beunders, G., van de Kamp, J., Vasudevan, P., Morton, J., Smets, K., Kleefstra, T., de Munnik, S.A., Schuurs-Hoeijmakers, J., Ceulemans, B., Zollino, M., et al. (2016). A detailed clinical analysis of 13 patients with AUTS2 syndrome further delineates the phenotypic spectrum and underscores the behavioural phenotype. J Med Genet 53, 523–532. 10.1136/jmedgenet-2015-103601.

7. Beunders, G., Voorhoeve, E., Golzio, C., Pardo, L.M., Rosenfeld, J.A., Talkowski, M.E., Simonic, I., Lionel, A.C., Vergult, S., Pyatt, R.E., et al. (2013). Exonic deletions in AUTS2 cause a syndromic form of intellectual disability and suggest a critical role for the C terminus. Am J Hum Genet 92, 210–220. 10.1016/j.ajhg.2012.12.011.

8. Elia, J., Gai, X., Xie, H.M., Perin, J.C., Geiger, E., Glessner, J.T., D’arcy, M., deBerardinis, R., Frackelton, E., Kim, C., et al. (2010). Rare structural variants found in attention-deficit hyperactivity disorder are preferentially associated with neurodevelopmental genes. Mol Psychiatry 15, 637–646. 10.1038/mp.2009.57.

9. Talkowski, M.E., Rosenfeld, J.A., Blumenthal, I., Pillalamarri, V., Chiang, C., Heilbut, A., Ernst, C., Hanscom, C., Rossin, E., Lindgren, A.M., et al. (2012). Sequencing chromosomal abnormalities reveals neurodevelopmental loci that confer risk across diagnostic boundaries. Cell 149, 525– 537. 10.1016/j.cell.2012.03.028.

10. Schumann, G., Coin, L.J., Lourdusamy, A., Charoen, P., Berger, K.H., Stacey, D., Desrivières, S., Aliev, F.A., Khan, A.A., Amin, N., et al. (2011). Genome-wide association and genetic functional studies identify autism susceptibility candidate 2 gene (AUTS2) in the regulation of alcohol consumption. Proc Natl Acad Sci U S A 108, 7119–7124. 10.1073/pnas.1017288108.

11. Chen, Y.-H., Liao, D.-L., Lai, C.-H., and Chen, C.-H. (2013). Genetic analysis of AUTS2 as a susceptibility gene of heroin dependence. Drug Alcohol Depend 128, 238–242. 10.1016/j.drugalcdep.2012.08.029.

12. Mefford, H.C., Muhle, H., Ostertag, P., von Spiczak, S., Buysse, K., Baker, C., Franke, A., Malafosse, A., Genton, P., Thomas, P., et al. (2010). Genome-wide copy number variation in epilepsy: novel susceptibility loci in idiopathic generalized and focal epilepsies. PLoS Genet 6, e1000962. 10.1371/journal.pgen.1000962.

13. Zhang, B., Xu, Y.-H., Wei, S.-G., Zhang, H.-B., Fu, D.-K., Feng, Z.-F., Guan, F.-L., Zhu, Y.-S., and Li, S.-B. (2014). Association study identifying a new susceptibility gene (AUTS2) for schizophrenia. Int J Mol Sci 15, 19406–19416. 10.3390/ijms151119406.

14. Ozsoy, F., Karakus, N.B., Yigit, S., and Kulu, M. (2020). Effect of AUTS2 gene rs6943555 variant in male patients with schizophrenia in a Turkish population. Gene 756, 144913. 10.1016/j.gene.2020.144913.

15. Girirajan, S., Brkanac, Z., Coe, B.P., Baker, C., Vives, L., Vu, T.H., Shafer, N., Bernier, R., Ferrero, G.B., Silengo, M., et al. (2011). Relative burden of large CNVs on a range of neurodevelopmental phenotypes. PLoS Genet 7, e1002334. 10.1371/journal.pgen.1002334.

16. Doust, C., Fontanillas, P., Eising, E., Gordon, S.D., Wang, Z., Alagöz, G., Molz, B., 23andMe Research Team, Quantitative Trait Working Group of the GenLang Consortium, Pourcain, B.S., et al. (2022). Discovery of 42 genome-wide significant loci associated with dyslexia. Nat Genet 54, 1621–1629. 10.1038/s41588-022-01192-y.

17. Biel, A., Castanza, A.S., Rutherford, R., Fair, S.R., Chifamba, L., Wester, J.C., Hester, M.E., and Hevner, R.F. (2022). AUTS2 Syndrome: Molecular Mechanisms and Model Systems. Front Mol Neurosci 15, 858582. 10.3389/fnmol.2022.858582.

18. Shimogori, T., Abe, A., Go, Y., Hashikawa, T., Kishi, N., Kikuchi, S.S., Kita, Y., Niimi, K., Nishibe, H., Okuno, M., et al. (2018). Digital gene atlas of neonate common marmoset brain. Neurosci Res 128, 1–13. 10.1016/j.neures.2017.10.009.

19. Kita, Y., Nishibe, H., Wang, Y., Hashikawa, T., Kikuchi, S.S. U M., Yoshida, A.C., Yoshida, C., Kawase, T., Ishii, S., et al. (2021). Cellular-resolution gene expression profiling in the neonatal marmoset brain reveals dynamic species- and region-specific differences. Proc Natl Acad Sci U S A 118, e2020125118. 10.1073/pnas.2020125118.

20. Lepagnol-Bestel, A.-M., Maussion, G., Boda, B., Cardona, A., Iwayama, Y., Delezoide, A.-L., Moalic, J.-M., Muller, D., Dean, B., Yoshikawa, T., et al. (2008). SLC25A12 expression is associated with neurite outgrowth and is upregulated in the prefrontal cortex of autistic subjects. Mol Psychiatry 13, 385–397. 10.1038/sj.mp.4002120.

21. Bedogni, F., Hodge, R.D., Nelson, B.R., Frederick, E.A., Shiba, N., Daza, R.A., and Hevner, R.F. (2010). Autism susceptibility candidate 2 (Auts2) encodes a nuclear protein expressed in developing brain regions implicated in autism neuropathology. Gene Expr Patterns 10, 9–15. 10.1016/j.gep.2009.11.005.

22. Yamashiro, K., Hori, K., Lai, E.S.K., Aoki, R., Shimaoka, K., Arimura, N., Egusa, S.F., Sakamoto, A., Abe, M., Sakimura, K., et al. (2020). AUTS2 Governs Cerebellar Development, Purkinje Cell Maturation, Motor Function and Social Communication. iScience 23, 101820. 10.1016/j.isci.2020.101820.

23. Besnard, A., and Leroy, F. (2022). Top-down regulation of motivated behaviors via lateral septum sub-circuits. Mol Psychiatry 27, 3119–3128. 10.1038/s41380-022-01599-3.

24. Burguière, E., Monteiro, P., Mallet, L., Feng, G., and Graybiel, A.M. (2015). Striatal circuits, habits, and implications for obsessive-compulsive disorder. Curr Opin Neurobiol 30, 59–65. 10.1016/j.conb.2014.08.008.

25. Josselyn, S.A., and Tonegawa, S. (2020). Memory engrams: Recalling the past and imagining the future. Science 367. 10.1126/science.aaw4325.

26. Russo, S.J., and Nestler, E.J. (2013). The brain reward circuitry in mood disorders. Nat Rev Neurosci 14, 609–625. 10.1038/nrn3381.

27. Dehaene, S., Cohen, L., Morais, J., and Kolinsky, R. (2015). Illiterate to literate: behavioural and cerebral changes induced by reading acquisition. Nat Rev Neurosci 16, 234–244. 10.1038/nrn3924.

28. Raslau, F.D., Mark, I.T., Klein, A.P., Ulmer, J.L., Mathews, V., and Mark, L.P. (2015). Memory part 2: the role of the medial temporal lobe. AJNR Am J Neuroradiol 36, 846–849. 10.3174/ajnr.A4169.

29. Hitti, F.L., and Siegelbaum, S.A. (2014). The hippocampal CA2 region is essential for social memory. Nature 508, 88–92. 10.1038/nature13028.

30. Middleton, S.J., and McHugh, T.J. (2020). CA2: A Highly Connected Intrahippocampal Relay. Annu Rev Neurosci 43, 55–72. 10.1146/annurev-neuro-080719-100343.

31. Johansen, J.P., Cain, C.K., Ostroff, L.E., and LeDoux, J.E. (2011). Molecular mechanisms of fear learning and memory. Cell 147, 509–524. 10.1016/j.cell.2011.10.009.

32. Smith, J.B., Lee, A.K., and Jackson, J. (2020). The claustrum. Curr Biol 30, R1401–R1406. 10.1016/j.cub.2020.09.069.

33. Crick, F.C., and Koch, C. (2005). What is the function of the claustrum? Philos Trans R Soc Lond B Biol Sci 360, 1271–1279. 10.1098/rstb.2005.1661.

34. Peng, H., Xie, P., Liu, L., Kuang, X., Wang, Y., Qu, L., Gong, H., Jiang, S., Li, A., Ruan, Z., et al. (2021). Morphological diversity of single neurons in molecularly defined cell types. Nature 598, 174–181. 10.1038/s41586-021-03941-1.

35. Livingstone, M.S., Rosen, G.D., Drislane, F.W., and Galaburda, A.M. (1991). Physiological and anatomical evidence for a magnocellular defect in developmental dyslexia. Proc Natl Acad Sci U S A 88, 7943–7947. 10.1073/pnas.88.18.7943.

36. Spiteri, S., and Crewther, D. (2021). Neural Mechanisms of Visual Motion Anomalies in Autism: A Two-Decade Update and Novel Aetiology. Front Neurosci 15, 756841. 10.3389/fnins.2021.756841.

37. Charpak, G., Dominik, W., and Zaganidis, N. (1989). Optical imaging of the spatial distribution of beta-particles emerging from surfaces. Proc Natl Acad Sci U S A 86, 1741–1745. 10.1073/pnas.86.6.1741.

38. O’Rahilly, R., and Müller, F. (1994). Neurulation in the normal human embryo. Ciba Found Symp 181, 70–82; discussion 82-89. 10.1002/9780470514559.ch5.

39. Silberberg, S.N., Taher, L., Lindtner, S., Sandberg, M., Nord, A.S., Vogt, D., Mckinsey, G.L., Hoch, R., Pattabiraman, K., Zhang, D., et al. (2016). Subpallial Enhancer Transgenic Lines: a Data and Tool Resource to Study Transcriptional Regulation of GABAergic Cell Fate. Neuron 92, 59–74. 10.1016/j.neuron.2016.09.027.

40. Nery, S., Fishell, G., and Corbin, J.G. (2002). The caudal ganglionic eminence is a source of distinct cortical and subcortical cell populations. Nat Neurosci 5, 1279–1287. 10.1038/nn971.

41. Lim, L., Mi, D., Llorca, A., and Marín, O. (2018). Development and Functional Diversification of Cortical Interneurons. Neuron 100, 294–313. 10.1016/j.neuron.2018.10.009.

42. Hu, J.S., Vogt, D., Lindtner, S., Sandberg, M., Silberberg, S.N., and Rubenstein, J.L.R. (2017). Coup-TF1 and Coup-TF2 control subtype and laminar identity of MGE-derived neocortical interneurons. Development 144, 2837–2851. 10.1242/dev.150664.

43. Bandler, R.C., Mayer, C., and Fishell, G. (2017). Cortical interneuron specification: the juncture of genes, time and geometry. Curr Opin Neurobiol 42, 17–24. 10.1016/j.conb.2016.10.003.

44. Xu, Q., Tam, M., and Anderson, S.A. (2008). Fate mapping Nkx2.1-lineage cells in the mouse telencephalon. J Comp Neurol 506, 16–29. 10.1002/cne.21529.

45. Lepagnol-Bestel, A.M., duchon, A., Viard, J., kvajo, M., Daudin, R., Khelfaoui, M., Haziza, S., Loe-Mie, Y., Aime, M., Suizu, F., et al. (2022). AUTS2 gene dosage affects synaptic AMPA receptors via a local dendritic spine AUTS2-TTC3-AKT-mTORC1 signaling dysfunction. bioRxiv.

46. Dölen, G., Malenka, R.C., Perlmutter, J.S., Brose, N., Frackowiak, R., Cuthbert, B.N., Diester, I., Mansuy, I., Kroker, K.S., Boeckers, T.M., et al. (2015). Pathophysiological Toolkit: Genes to Circuits. In Translational Neuroscience: Toward New Therapies, K. Nikolich and S. E. Hyman, eds. (MIT Press).

47. Hori, K., Nagai, T., Shan, W., Sakamoto, A., Abe, M., Yamazaki, M., Sakimura, K., Yamada, K., and Hoshino, M. (2015). Heterozygous Disruption of Autism susceptibility candidate 2 Causes Impaired Emotional Control and Cognitive Memory. PLoS One 10, e0145979. 10.1371/journal.pone.0145979.

48. Hori, K., Yamashiro, K., Nagai, T., Shan, W., Egusa, S.F., Shimaoka, K., Kuniishi, H., Sekiguchi, M., Go, Y., Tatsumoto, S., et al. (2020). AUTS2 Regulation of Synapses for Proper Synaptic Inputs and Social Communication. iScience 23, 101183. 10.1016/j.isci.2020.101183.

49. Li, J., Sun, X., You, Y., Li, Q., Wei, C., Zhao, L., Sun, M., Meng, H., Zhang, T., Yue, W., et al. (2022). Auts2 deletion involves in DG hypoplasia and social recognition deficit: The developmental and neural circuit mechanisms. Sci Adv 8, eabk1238. 10.1126/sciadv.abk1238.

50. Engmann, O., Labonté, B., Mitchell, A., Bashtrykov, P., Calipari, E.S., Rosenbluh, C., Loh, Y.-H.E., Walker, D.M., Burek, D., Hamilton, P.J., et al. (2017). Cocaine-Induced Chromatin Modifications Associate With Increased Expression and Three-Dimensional Looping of Auts2. Biol Psychiatry 82, 794–805. 10.1016/j.biopsych.2017.04.013.

51. Misra, V. (2014). The social brain network and autism. Ann Neurosci 21, 69–73. 10.5214/ans.0972.7531.210208.

52. Corbin, J.G., Nery, S., and Fishell, G. (2001). Telencephalic cells take a tangent: non-radial migration in the mammalian forebrain. Nat Neurosci 4 Suppl, 1177–1182. 10.1038/nn749.

53. Loomba S, Straehle J, Gangadharan V, Heike N, Khalifa A, Motta A, Ju N, Sievers M, Gempt J, Meyer HS, Helmstaedter M. (2022). Connectomic comparison of mouse and human cortex. Science. Jul 8;377(6602):eabo0924.

54. Krienen FM, Goldman M, Zhang Q, C H Del Rosario R, Florio M, Machold R, Saunders A, Levandowski K, Zaniewski H, Schuman B, Wu C, Lutservitz A, Mullally CD, Reed N, Bien E, Bortolin L, Fernandez-Otero M, Lin JD, Wysoker A, Nemesh J, Kulp D, Burns M, Tkachev V, Smith R, Walsh CA, Dimidschstein J, Rudy B, S Kean L, Berretta S, Fishell G, Feng G, McCarroll SA.(2020). Innovations present in the primate interneuron repertoire. Nature.Oct;586(7828):262–269.

